# Gene expression genetics of the striatum of Diversity Outbred mice

**DOI:** 10.1101/2023.05.11.540390

**Authors:** Vivek M. Philip, Hao He, Michael C. Saul, Price E. Dickson, Jason A. Bubier, Elissa J. Chesler

## Abstract

Brain transcriptional variation is a heritable trait that mediates complex behaviors, including addiction. Expression quantitative trait locus (eQTL) mapping reveals genomic regions harboring genetic variants that influence transcript abundance. In this study, we profiled transcript abundance in the striatum of 386 Diversity Outbred (J:DO) mice of both sexes using RNA-Seq. All mice were characterized using a behavioral battery of widely-used exploratory and risk-taking assays prior to transcriptional profiling. We performed eQTL mapping, incorporated the results into a browser-based eQTL viewer, and deposited co-expression network members in GeneWeaver. The eQTL viewer allows researchers to query specific genes to obtain allelic effect plots, analyze SNP associations, assess gene expression correlations, and apply mediation analysis to evaluate whether the regulatory variant is acting through the expression of another gene. GeneWeaver allows multi-species comparison of gene sets using statistical and combinatorial tools. This data resource allows users to find genetic variants that regulate differentially expressed transcripts and place them in the context of other studies of striatal gene expression and function in addiction-related behavior.

## Background & Summary

Substance use disorder is a highly heritable trait involving variation in neural circuitry underlying motivated behavior and behavioral inhibition. Characterization of addiction-related brain regions in genetically diverse mice can lead to the discovery of molecular mechanisms of addiction-related behaviors. These mechanisms can in turn, aid in connecting genetic, genomic and behavioral variation within and across species through shared target genes^1^.

Drug-induced transcriptional changes in the corticostriatal system have been reported in rodent models^2,3^ and substance dependent individuals^4,5^. Additionally, behavioral correlates of substance use disorder, namely impulsivity and incentive sensitization, all involve corticostriatal circuitry^6-10^; however, the molecular mechanisms underlying these relationships are unknown. The striatum plays a central role in addiction-related behavior ^11,12^ and influences behaviors (e.g., sensation seeking) that predict the development of substance use disorders ^13-16^. It receives inputs from diverse brain regions, including the midbrain, prefrontal cortex, and thalamus, and plays a fundamental role in goal-directed actions and habits ^11^. The dopamine projections from the ventral tegmental area to the nucleus accumbens and prefrontal cortex are at the heart of this reward circuit, and their importance to drug reward is well established^17^. Neuroimaging studies of people with a history of cocaine use disorders and rodent studies have indicated that addiction is a circuit-level disorder involving several functionally inter-connected brain regions^18,19^. Identifying drug-induced changes in gene expression^20,21^ and resulting neural plasticity^22,23^ in addiction-relevant neurocircuits can reveal underlying sources of addiction risk and resilience.

Gene expression quantitative trait locus (eQTL) and systems genetic analyses facilitate the identification of genes and variants associated with complex traits, including those related to addiction.^24^ Such data allow the reduction of large numbers of positional candidate genes and variants implicated in quantitative trait locus (QTL) for behavioral traits. These data are also useful for discovering transcripts significantly correlated with behavior and uncovering transcriptional co-expression networks to identify the biomolecular mechanisms underlying complex traits^24^. Researchers can also use eQTL data to identify genetic variants regulating differentially expressed genes, such as those discovered in drug exposure studies. This data can be used with data from other model organisms^25^ to identify convergent evidence for biological mechanisms of addiction across species^26,27^. Model organism eQTLs can also be related to convergent findings in humans to prioritize genome-wide association study (GWAS) results and to contextualize the role of the identified variant^1^.

To ensure variation in nearly every gene in the genome and to increase the precision of QTL mapping, the Diversity Outbred^28-30^ (J:DO) mice were developed as an advanced intercross of the eight-way hybrid Collaborative Cross (CC) population^31,32^. Within the J:DO population, there are over 45 million single-nucleotide polymorphisms (SNPs) and millions of insertions and deletions segregating^12,33,34^. This high genetic diversity results in high phenotypic and transcriptomic variation ^35,36^, enabling the discovery of genes and variants associated with behaviors.

Transcription regulatory variation is often context specific. Studies of transcriptional variation in J:DO mice have revealed precise genetic variation affecting proteomes and cellular transcriptional states in addition to bulk transcriptomics of tissues relevant to metabolism in health and disease^37-39^.

To construct a versatile reference data resource for addiction genetics, we performed a series of addiction-relevant exploratory and risk-taking behavioral assays^35,40^, and then profiled transcript abundance in striatum using RNA-Seq on 368 drug naïve J:DO mice. Data from this study are delivered in a platform that allows for the identification of eQTL effects, analysis of local SNP associations, assessment of gene expression correlations, and application of mediation analysis. This provides a resource for genetic studies of transcriptional diversity in the striatum of drug naïve mice. Combined with behavioral phenotyping, this resource enables the prioritization of behavioral QTL positional candidates by incorporating evidence from strong cis-eQTL effects and their underlying allelic patterns. Furthermore, behavioral QTLs can be subjected to local SNP association analysis followed by prioritization of positional candidates where the SNP strain distribution pattern of positional candidates matches the allelic effects of interest. Positional candidates can then be queried in this resource for the presence or absence of strong cis-eQTLs. Finally, the data can be used in global analyses of the relationship of trait-relevant variation across species, using increasingly sophisticated approaches for leveraging model organism data to predict, model, and explain polygenic risk for human disease ^41-43^.

## Methods

### Mice

416 J:DO mice (strain #:009376) from both sexes, spanning generations G21, G22, and G23, were purchased from The Jackson Laboratory. The mice were housed in an elevated barrier pathogen- and opportunistic-free animal room (Health report available at: https://www.jax.org/-/media/jaxweb/health-reports/g200.pdf?la=en&hash=7AD522E82FA7C6D614A11EFB82547476157F00E1) before being transferred at weaning to an intermediate barrier specific pathogen-free room (https://www.jax.org/-/media/jaxweb/health-reports/g3b.pdf?la=en&hash=914216EE4F44ADC1585F1EF219CC7F631F881773). Mice were individually housed under (12:12) light/dark cycle and allowed *ad libitum* access to standard rodent chow [sterilized NIH31 5K52 6% fat chow (LabDiet/PMI Nutrition, St. Louis, MO)] and acidified water (pH 2.5–3.0) supplemented with vitamin K. Mice cages contained a pine-shaving bedding (Hancock Lumber) and environmental enrichment consisting of a nestlet and a Shepard’s Shack. The mice were identified by ear notching at weaning and moved between cages and testing using metal forceps.

### Behavioral Phenotyping

At three to six months of age, mice were phenotyped four separate times with a different assay on each day, Monday to Thursday (open field, light-dark, hole-board, and novelty place preference)^44^ and euthanized on Friday in batches of 16-24 by decapitation. Phenotyping protocols are available at https://www.addiction-neurogenetics.org/data-and-resources/. The Jackson Laboratory (JAX) follows husbandry practices in accordance with the American Association for the Accreditation of Laboratory Animal Care (AAALAC), and all work was done with the approval of the JAX Institutional Animal Care and Use Committee (Approval #10007).

### Dissections

Testing and euthanasia were consistently performed between 8 AM to 12 PM to control for circadian effects. All surgical instruments were cleaned with RNAase Away (ThermoFischer Scientific, Waltham, MA) prior to use and between samples. Whole intact brains were removed, hemisected, and incubated in RNAlater (ThermoFischer Scientific, Waltham, MA) for 8-14 minutes. Then, under a dissection microscope, the striatum, hippocampus, and prefrontal cortex were removed and soaked for 24 hours in RNAlater at room temperature before being stored at -80°C until processing.

### RNA Isolation and Sequencing

The striatum was homogenized, and total RNA was isolated using a TRIzol Plus kit (Life Technologies, City, State) with on-the-column DNase digestion according to the manufacturer’s instructions. The quality of the isolated RNA was assessed using an RNA 6000 Nano LabChip using an Agilent 2100 Bioanalyzer instrument [RRID:SCR_019389 (Agilent Technologies, Santa Clara, CA)] and a NanoDrop spectrophotometer [RRID:SCR_018042 (ThermoFisher Scientific, Wilmington, DE)]. The External RNA Controls Consortium spike-in (ERCC, Ambion, Austin, TX) was added to the samples to allow for normalization in accordance with the core facility’s standard operating procedure but was not used in our downstream analyses. An RNA-Seq library was prepared using the KAPA Stranded RNA-Seq Kit with RiboErase (Kappa Biosystems, City, State). Libraries were then pooled and sequenced at The Jackson Laboratory using a 100 bp paired-end process on a HiSeq 2500 (Illumina) sequencing system (RRID:SCR_016383) targeting 40 million read pairs per sample. Sequencing achieved a median read depth of 65.7 million read pairs per sample (range: 31.4 million to 117.4 million reads).

### Sequencing Analysis

Raw read data were demultiplexed and converted to FASTQ files. Paired-end FASTQ files from multiple lanes were concatenated together prior to alignment. All paired-end FASTQ files were aligned to the *Mus musculus* GRCm38 reference (GenBank accession number: GCA_000001635.2) with Ensembl v94 (October 2018) annotation using STAR (v2.6.1c) (RRID:SCR_004463) set to produce both genome and transcriptome Binary Alignment Map (BAM) files. STAR was used with default options ensuring that these defaults allowed for a maximum of 10 multi-mapped reads and a maximum of 10 mismatches. Reads exceeding these criteria were excluded from further downstream processing. Expression estimation was performed using the RSEM package (v1.3.0) (RRID:SCR_013027) with --estimate-rspd using the transcriptome BAM files obtained following alignments from STAR. RSEM expected counts per transcript were used for downstream analysis. Expression estimate data were imported into R v3.5.1 (RRID:SCR_001905) using tximport v 1.10.1 (RRID:SCR_016752). Data were TMM-normalized with edgeR v3.24.3 (RRID:SCR_012802) and log-transformed to stabilize variance using voom+limma in limma v3.38.3 (RRID:SCR_010943). We used the biomaRt R package v2.38.0 (RRID:SCR_019214) to annotate the data using the v94 Ensembl archive (oct2018.archive.ensembl.org). Using X and Y chromosome gene expression, we discovered that some samples had sex chromosome aneuploidies (X0 females and partial XXY males), a previously documented phenomenon among J:DO mice ^45^. These samples were excluded from downstream analyses. Additionally, we discovered that some samples included choroid plexus contamination. We remediated this contamination by taking the residuals of expression regressed on the log-mean expression of the genes klotho (*Kl*, ENSMUSG00000058488) and transthyretin (*Ttr*, ENSMUSG00000061808), which are unambiguous markers for the choroid plexus ^46^.

### Genotyping, Haplotype Reconstruction and Sample QC

Genotyping was performed on tail biopsies by Neogene Genomics (Lincoln, NE) using the Mouse Universal Genotyping Array (GigaMUGA)^47^ consisting of 143,259 markers. Based on published genotype QC workflows^48^, 110,524 markers and 386 mice were retained for further analysis. ^48^. Genotypes were converted to founder strain-haplotype reconstructions using R/qtl2^49^(qtl2_0.21-1, http://kbroman.org/qtl2) (RRID:SCR_018181).

### Expression QTL mapping

Prior to eQTL mapping, gene expression counts were obtained by summing expected counts over all transcripts for a given gene. Expression for eQTL analysis was adjusted for choroid plexus contamination by regressing the log-mean of *Kl* and *Ttr* as additive covariates. eQTL mapping was performed on regression residuals of 17,248 genes using the R/qtl2 package and the founder haplotype regression method. To correct for population structure, kinship matrices were computed with the Leave One Chromosome Out (LOCO) option for kinship correction (http://kbroman.org/qtl2)^50^. Additive covariates of sex and J:DO generation were used in the eQTL mapping model. Specifically, for each gene, the following linear model was fit,

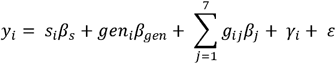

where *y*_*i*_ is the gene expression abundance of the *i*^*th*^ animal, *s*_*i*_ is the sex of animal *i, β*_*s*_ is the effect of sex, *gen*_*i*_ is the generation of animal *i, β*_*gen*_ is the effect of generation, *g*_*ij*_ is the founder probability for founder allele *j* in animal *i, β*_j_ is the genotype coefficient, *γ*_*i*_ and is the random effect representing the polygenic influence of animal *i* as modeled by a kinship matrix. eQTL was categorized as either cis or trans, where cis is defined as eQTLs within +/-2MB of the transcription start site (TSS) of the gene, and trans are eQTLs further away.

### Creation of the eQTL Viewer Object

The results of the eQTL analysis, along with expression estimates, genotypes, and covariates, are encapsulated in an RData object. RData objects are designed for use in R and contain all the objects necessary for reproducing an analysis. We followed the instructions provided by the developers of the eQTL viewer^51,52^. Specifically, we created the following elements: kinship, map, genotype probabilities (genoprobs), markers, and a dataset object that contains information on the gene annotations, covariates, expression data, and sample annotations.

### WGCNA Analysis

RNA-Seq data was analyzed with WGCNA (RRID:SCR_003302) ^53^. A soft thresholding power of 3 was selected using the WGCNA scale-free topology R^2^ threshold of 0.9 with a signed network with a minimum module size of 30. The correlation calculation utilized was bicor, and modules used numeric labels instead of colors.

### Paraclique analysis

RNA-Seq data was analyzed with paraclique^54^ using a bicor with a correlation coefficient threshold of |0.5| (unsigned), minimum seed clique size of 5, minimum finished paraclique size of 10, proportional glom factor of 0.2 for paraclique construction.

### Genesets for Analysis in GeneWeaver

Set of genes representing the J:DO striatum eQTL, the WGCNA modules and the paracliques were deposited in GeneWeaver (RRID:SCR_009202)^55^ are accession numbers are found in Supplemental Table 1. In addition the eQTLs have been separated into cis and trans sets and are also presented as sets per chromosome.

## Data Records

### Primary Sequence Data

Primary raw paired end RNA-Seq data files (FASTQ formatted) from 416 J:DO mice were submitted to the Sequence Read Archive (SRA) and are available with the GEO ID (GSE162732).

### Primary Genotyping Data

Raw data has been deposited at the Diversity Outbred Database https://www.jax.org/research-and-faculty/genetic-diversity-initiative/tools-data/diversity-outbred-database (RRID:SCR_018180)

### Primary Phenotyping Data

Phenotyping data has been deposited at the Mouse Phenome Database (RRID:SCR_003212) under project CSNA03.

### QTL Viewer Repository

QTL Viewer is an interactive web-based analysis tool allowing users to replicate the analyses reported for a study (**Figure 1**). The tool with the data set described here is available at https://qtlviewer.jax.org/viewer/CheslerStriatum. It includes the ability to search various subsets of data from a study, such as phenotypes or expression data, and then visualize data with profile, correlation, LOD, effect, mediation, and SNP association plots (**Figure 2**). Detailed information about the structure of the QTL viewer objects is available at https://github.com/churchill-lab/qtl-viewer/blob/master/docs/QTLViewerDataStructures.md. A complementary dataset from the hippocampus previous described in Skeelly et.al^56^ is already available at https://churchilllab.jax.org/qtlviewer/DO/hippocampus

**Figure 1.**
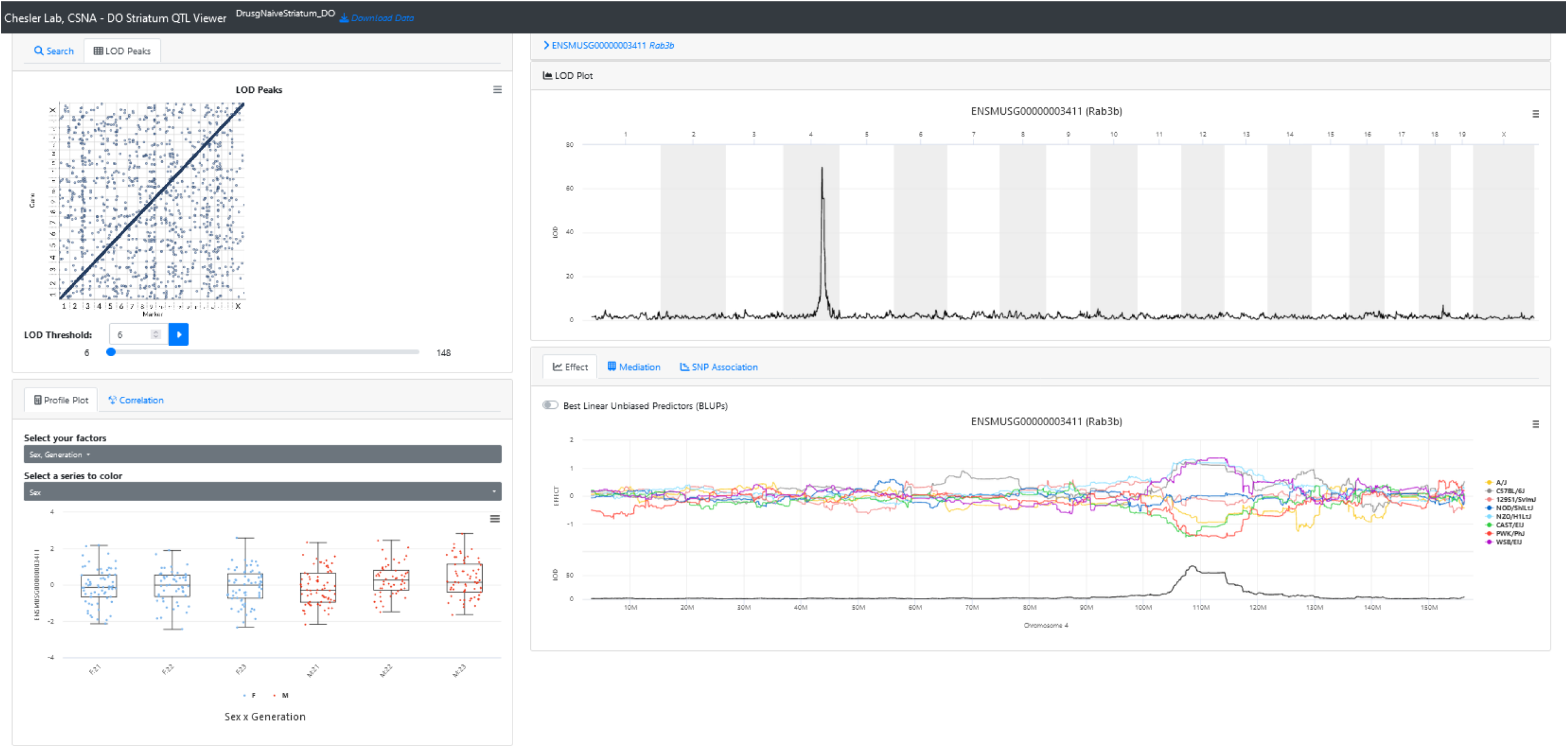
Screenshot of a query for the gene “*Rab3b*” in QTL Viewer. QTL results for *Rab3b* expression in the Diversity Outbred (J:DO) mice striatum. Metadata related to the J:DO generation and sex is displayed, and genes co-expressed with the selected gene can be accessed from the correlation tab. The allele effect plots, SNP association mapping, and mediation analysis can also be performed and viewed from the page.

**Figuer 2.**
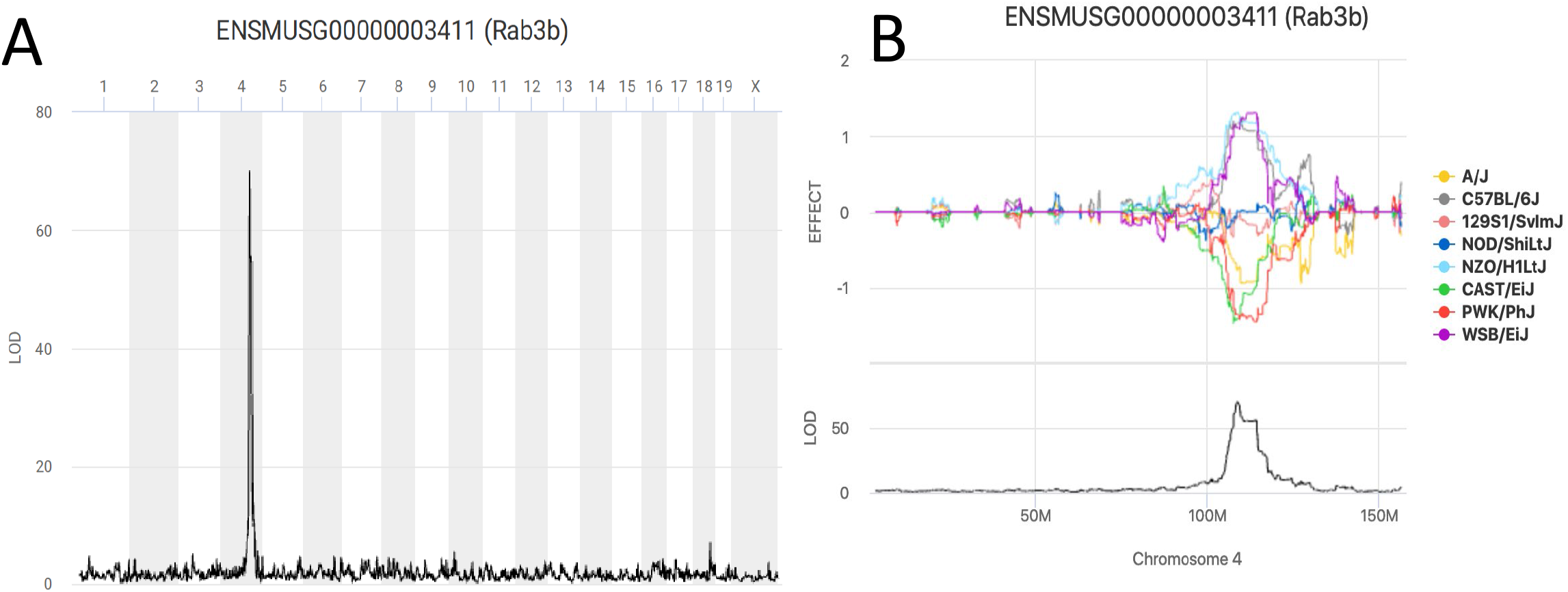
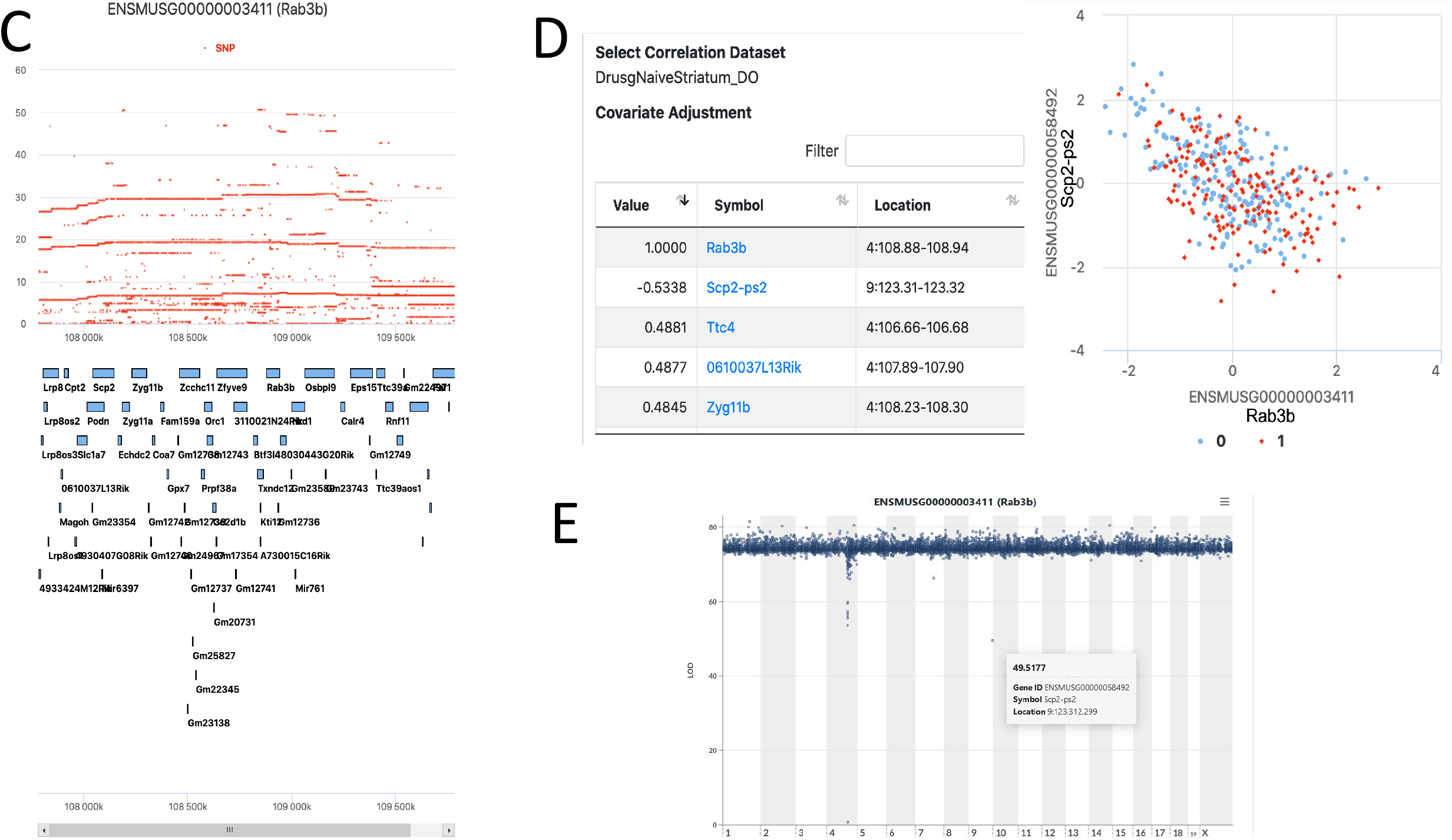
The *Rab3b* cis-eQTL data retrieved in QTL Viewer. **A**. A genome scan for *Rab3b* expression in the Diversity Outbred (J:DO) mice striatum identifies a strong (LOD > 65) cis-eQTL on chromosome 4. **B**. The allele effect plot for the haplotypes of the J:DO that regulate the expression of *Rab3b*. There are strong effects of WSB/EiJ, NZO/HILtJ, and C57BL/6J on the expression in one direction and 129S1/SvJ, CAST/EiJ, and A/J in the opposite direction. **C**. SNP association mapping within the QTL peak interval displaying all the SNPs that drive the QTL. Most of the highest score SNPs are around *Rab3b*, as expected for a gene regulated in cis. **D**. The genes that are negatively (*Scp2*-ps2) or positively (*Ttc4, 0610037L13Rik, Zyg11b*) correlated with *Rab3b* expression can be displayed. A scatter plot can be generated for each gene with the gene of interest. **E**. Mediation analysis can be performed on the data set to identify candidate causal mediators. This analysis retests the QTL effect at the locus of interest, iteratively conditioned on candidate mediators. Here the SNP in Scp2-ps2 creates the greatest LOD drop.

### QTL Viewer RData object

The primary data record associated with this study is qtlviewer_DO_Striatum_02102020.Rdata. This RData object contains the following:

- genoprobs - the genotype probabilities
- K - the kinship matrix created using the leave one chromosome out (loco) method
- map - list of one element per chromosome, with the genomic position of each marker
- markers - marker names and positions
- dataset.DO_Striatum_416 –
- annot.mrna - annotations of the mRNA data
  - annot.samples - sample annotations
  - covar.info - specific information about the covariates
  - covar.matrix - matrix of covariates data, samples (rows) x covariates (columns)
  - data - expression data, samples (rows) x mRNA (columns). This matrix is used in the eQTL mapping analysis.
  - datatype - type of data set, either mRNA or protein
  - display.name - simple display name for the viewer
  - ensembl.version - version of Ensembl used to annotate locations
  - lod.peaks - LOD peaks over a certain threshold, set to >7 in this dataset

### Technical Validation

#### Blinding and Randomization

In a population genetics study, mouse genotypes are collected randomly. The J:DO population is bred using a pseudorandom mating scheme, and test mice are obtained from several breeding cohorts. Experimenters are unaware of mouse genotypes and their relationship to gene expression. Coat color diversity in this population may create some experimenter bias in conventional mouse populations, but coat color is rarely a predictor of behavior in J:DO mice. In the ideal genetic population, genotypes are fully randomized, and individuals are all genetically equidistant. Because this is not the case, genetic mapping analyses include a relationship matrix, a structured covariance matrix that estimates the relations among individuals based on genotype similarity. Mice of both sexes are counterbalanced across test runs, with each run containing either male or female mice to avoid pheromonal effects on behavior.

### RNA quality

RNA quality includes three primary components: integrity, purity, and concentration. RNA integrity was determined by the Agilent Bioanalyzer 2100 using an RNA Integrity Number (RIN). This metric uses the ratio of 28S:18S rRNA to rate RNA quality on a scale of 1-10. The median RIN was 9.1 (range: 7.9-9.9). RNA purity and concentration were determined using a NanoDrop spectrophotometer. Using this method, RNA concentration is determined using absorbance at 260 nm, while purity is determined using the ratio of absorbance at 260 nm to 280 nm (A260:A280) and the ratio of absorbance at 260 nm to 230 nm (A260:A230). The median concentration was 74.0 ng/μL (range: 9.4-186.3 ng/μL), the median A260:A280 ratio was 2.00 (range:1.67-2.06), and the median A260:A230 ratio was 2.02 (range: 0.98-3.23). These values indicated that the RNA quality was sufficient for RNA-Seq analysis.

### Heritability of transcript variation

Based on variance accounted for by genotypes across the genome, and an additive covariant of generation, transcript abundance has a median heritability of 0.229, which is about two times the observed median heritability observed initially in the BXD^24^ Cis-eQTLs were highly detectable across the entire genome, as a diagonal band (seen in Figure 1). Trans-eQTLs were independent of each other in the genetically unstructured and large population as would be expected. UNC506203, on chr 1 and 40.21 Mbp and is the peak marker for the most number (42) of cis- and trans-eQTL. The largest interval between markers (13.852282 Mbp) that have no eQTL is on the X chromosome. This is all consistent with technically valid eQTL mapping.

### Usage Notes

There are four means by which the processed drug naïve striatum gene expression datasets can be used. First, users can download the entire processed dataset in the RData format. Once downloaded, it is readily readable in the R programming environment. Second, users can access the data at https://qtlviewer.jax.org/viewer/CheslerStriatum and proceed with the eQTL profile by entering their specific gene of interest in the search text box. Thirdly, users can access these data using the QTL viewer API interface (https://github.com/churchill-lab/qtl2api). Finally users can download sets of genes derived from the eQTL data by various methods such as WGCNA and paraclique, from the online data repository and suite of tools GeneWeaver.org.

## Code Availability

The R scripts and the package versions used for the eQTL analysis and the creation of the QTL viewer RData object are available at https://github.com/TheJacksonLaboratory/CSNA

## Acknowledgments

Major support for this work was from the National Institute on Drug Abuse of the National Institutes of Health (NIH RO1 DA037927; NIH P50 DA 039841, NIH U01 DA043809). We thank Tyler Roy and Troy Wilcox for performing the striatum dissections. We thank Gary Churchill and Matthew Vincent for their help with the QTL Viewer. This research was also supported by the Genome and Single Cell Technologies and Computational Sciences Shared Resources of the JAX Cancer Center (P30 CA034196). Finally, we gratefully acknowledge the contribution of Heidi Munger and Genome Technologies at The Jackson Laboratory for expert assistance with this publication.

## Author contributions

Vivek M. Philip-Wrote the manuscript and performed mapping studies.

Hao He – Prepared the genotype probabilities matrix and participated in QTL mapping.

Michael C. Saul – Performed the transcriptomic alignments, choroid plexus cleaning, QTL mapping and heritability calculations. Participated in the preparation of the manuscript.

Price E. Dickson – Oversaw the behavioral testing of the mice and participated in the preparation of the manuscript.

Jason A. Bubier. Coordinated procurement of, scheduling, genotyping, and dissecting of Diversity Outbreed mice. Participated in the preparation of the manuscript

Elissa J. Chesler. Conceived the project, secured funding, and oversaw the design and execution of the research and the manuscript.

## Competing interests

There are no competing interests to declare from authors of this manuscript.

## Figure Legends

**Supplemental Table 1: Table of J:DO striatum Gene Sets that are in GeneWeaver.**

## Notes

### Competing Interest Statement

The authors have declared no competing interest.

## References

1 Reynolds, T. et al. Interpretation of psychiatric genome-wide association studies with multispecies heterogeneous functional genomic data integration. Neuropsychopharmacology 46, 86–97 (2021). https://doi.org:10.1038/s41386-020-00795-5

2 Walker, D. M. et al. Cocaine Self-administration Alters Transcriptome-wide Responses in the Brain’s Reward Circuitry. Biol Psychiatry 84, 867–880 (2018). https://doi.org:10.1016/j.biopsych.2018.04.009

3 Huggett, S. B. et al. Genes identified in rodent studies of alcohol intake are enriched for heritability of human substance use. Alcohol Clin Exp Res (2021). https://doi.org:10.1111/acer.14738

4 Ribeiro, E. A. et al. Gene Network Dysregulation in Dorsolateral Prefrontal Cortex Neurons of Humans with Cocaine Use Disorder. Sci Rep 7, 5412 (2017). https://doi.org:10.1038/s41598-017-05720-3

5 Huggett, S. B. & Stallings, M. C. Genetic Architecture and Molecular Neuropathology of Human Cocaine Addiction. The Journal of neuroscience : the official journal of the Society for Neuroscience 40, 5300–5313 (2020). https://doi.org:10.1523/JNEUROSCI.2879-19.2020

6 Jupp, B. & Dalley, J. W. Convergent pharmacological mechanisms in impulsivity and addiction: insights from rodent models. Br J Pharmacol 171, 4729–4766 (2014). https://doi.org:10.1111/bph.12787

7 Jentsch, J. D. & Taylor, J. R. Impulsivity resulting from frontostriatal dysfunction in drug abuse: implications for the control of behavior by reward-related stimuli. Psychopharmacology (Berl) 146, 373–390 (1999). https://doi.org:10.1007/pl00005483

8 Meyer, P. J., King, C. P. & Ferrario, C. R. Motivational Processes Underlying Substance Abuse Disorder. Curr Top Behav Neurosci 27, 473–506 (2016). https://doi.org:10.1007/7854_2015_391

9 Kalivas, P. W., Pierce, R. C., Cornish, J. & Sorg, B. A. A role for sensitization in craving and relapse in cocaine addiction. J Psychopharmacol 12, 49–53 (1998). https://doi.org:10.1177/026988119801200107

10 Wolf, M. E. The Bermuda Triangle of cocaine-induced neuroadaptations. Trends Neurosci 33, 391–398 (2010). https://doi.org:10.1016/j.tins.2010.06.003

11 Everitt, B. J. & Robbins, T. W. Neural systems of reinforcement for drug addiction: from actions to habits to compulsion. Nature neuroscience 8, 1481–1489 (2005). https://doi.org:10.1038/nn1579

12 Belin, D. & Everitt, B. J. Cocaine seeking habits depend upon dopamine-dependent serial connectivity linking the ventral with the dorsal striatum. Neuron 57, 432–441 (2008). https://doi.org:10.1016/j.neuron.2007.12.019

13 Lind, N. M. et al. Behavioral response to novelty correlates with dopamine receptor availability in striatum of Gottingen minipigs. Behavioural brain research 164, 172–177 (2005). https://doi.org:10.1016/j.bbr.2005.06.008

14 Wittmann, B. C., Daw, N. D., Seymour, B. & Dolan, R. J. Striatal activity underlies novelty-based choice in humans. Neuron 58, 967–973 (2008). https://doi.org:10.1016/j.neuron.2008.04.027

15 Hooks, M. S. et al. Individual locomotor response to novelty predicts selective alterations in D1 and D2 receptors and mRNAs. The Journal of neuroscience : the official journal of the Society for Neuroscience 14, 6144–6152 (1994).

16 Gjedde, A., Kumakura, Y., Cumming, P., Linnet, J. & Moller, A. Inverted-U-shaped correlation between dopamine receptor availability in striatum and sensation seeking. Proceedings of the National Academy of Sciences of the United States of America 107, 3870–3875 (2010). https://doi.org:10.1073/pnas.0912319107

17 Cooper, S., Robison, A. J. & Mazei-Robison, M. S. Reward Circuitry in Addiction. Neurotherapeutics 14, 687–697 (2017). https://doi.org:10.1007/s13311-017-0525-z

18 Nielsen, G. [We are not proud enough]. Sygeplejersken 90, 18–20 (1990).

19 Luscher, C. The Emergence of a Circuit Model for Addiction. Annu Rev Neurosci 39, 257–276 (2016). https://doi.org:10.1146/annurev-neuro-070815-013920

20 James, M. H. Mimicking Human Drug Consumption Patterns in Rat Engages Corticostriatal Circuitry. Neuroscience 442, 311–313 (2020). https://doi.org:10.1016/j.neuroscience.2020.06.012

21 Sadri-Vakili, G. Cocaine triggers epigenetic alterations in the corticostriatal circuit. Brain Res 1628, 50–59 (2015). https://doi.org:10.1016/j.brainres.2014.09.069

22 Bobadilla, A. C. et al. Corticostriatal plasticity, neuronal ensembles, and regulation of drug-seeking behavior. Prog Brain Res 235, 93–112 (2017). https://doi.org:10.1016/bs.pbr.2017.07.013

23 Wall, N. R. et al. Complementary Genetic Targeting and Monosynaptic Input Mapping Reveal Recruitment and Refinement of Distributed Corticostriatal Ensembles by Cocaine. Neuron 104, 916–930 e915 (2019). https://doi.org:10.1016/j.neuron.2019.10.032

24 Chesler, E. J. et al. Complex trait analysis of gene expression uncovers polygenic and pleiotropic networks that modulate nervous system function. Nat Genet 37, 233–242 (2005). https://doi.org:10.1038/ng1518

25 Munro, D. et al. The regulatory landscape of multiple brain regions in outbred heterogeneous stock rats. Nucleic Acids Res 50, 10882–10895 (2022). https://doi.org:10.1093/nar/gkac912

26 Bubier, J. A. et al. Cross-Species Integrative Functional Genomics in GeneWeaver Reveals a Role for Pafah1b1 in Altered Response to Alcohol. Front Behav Neurosci 10, 1 (2016). https://doi.org:10.3389/fnbeh.2016.00001

27 Bubier, J. A. et al. Identification of a QTL in Mus musculus for alcohol preference, withdrawal, and Ap3m2 expression using integrative functional genomics and precision genetics. Genetics 197, 1377–1393 (2014). https://doi.org:10.1534/genetics.114.166165

28 Svenson, K. L. et al. High-resolution genetic mapping using the Mouse Diversity outbred population. Genetics 190, 437–447 (2012). https://doi.org:10.1534/genetics.111.132597 190/2/437 [pii]

29 Chesler, E. J. Out of the bottleneck: the Diversity Outcross and Collaborative Cross mouse populations in behavioral genetics research. Mamm Genome 25, 3–11 (2014). https://doi.org:10.1007/s00335-013-9492-9

30 Churchill, G. A., Gatti, D. M., Munger, S. C. & Svenson, K. L. The Diversity Outbred mouse population. Mamm Genome 23, 713–718 (2012). https://doi.org:10.1007/s00335-012-9414-2

31 Chesler, E. J. et al. The Collaborative Cross at Oak Ridge National Laboratory: developing a powerful resource for systems genetics. Mamm Genome 19, 382–389 (2008). https://doi.org:10.1007/s00335-008-9135-8

32 Churchill, G. A. et al. The Collaborative Cross, a community resource for the genetic analysis of complex traits. Nat Genet 36, 1133–1137 (2004). https://doi.org:ng1104-1133 x[pii] 10.1038/ng1104-1133

33 Ferraj, A. et al. Resolution of structural variation in diverse mouse genomes reveals chromatin remodeling due to transposable elements. bioRxiv, 2022.2009.2026.509577 (2022). https://doi.org:10.1101/2022.09.26.509577

34 Threadgill, D. W. & Churchill, G. A. Ten years of the Collaborative Cross. Genetics 190, 291–294 (2012). https://doi.org:10.1534/genetics.111.138032

35 Logan, R. W. et al. High-precision genetic mapping of behavioral traits in the diversity outbred mouse population. Genes Brain Behav 12, 424–437 (2013). https://doi.org:10.1111/gbb.12029

36 Philip, V. M. et al. Genetic analysis in the Collaborative Cross breeding population. Genome Res 21, 1223–1238 (2011). https://doi.org:10.1101/gr.113886.110 gr.113886.110 [pii]

37 Chick, J. M. et al. Defining the consequences of genetic variation on a proteome-wide scale. Nature 534, 500–505 (2016). https://doi.org:10.1038/nature18270

38 Skelly, D. A. et al. Mapping the Effects of Genetic Variation on Chromatin State and Gene Expression Reveals Loci That Control Ground State Pluripotency. Cell Stem Cell 27, 459–469 e458 (2020). https://doi.org:10.1016/j.stem.2020.07.005

39 Keele, G. R. et al. Regulation of protein abundance in genetically diverse mouse populations. Cell Genom 1 (2021). https://doi.org:10.1016/j.xgen.2021.100003

40 Recla, J. M. et al. Precise genetic mapping and integrative bioinformatics in Diversity Outbred mice reveals Hydin as a novel pain gene. Mamm Genome (2014). https://doi.org:10.1007/s00335-014-9508-0

41 Palmer, R. H. C. et al. Multi-omic and multi-species meta-analyses of nicotine consumption. Transl Psychiatry 11, 98 (2021). https://doi.org:10.1038/s41398-021-01231-y

42 Huggett, S. B., Bubier, J. A., Chesler, E. J. & Palmer, R. H. C. Do gene expression findings from mouse models of cocaine use recapitulate human cocaine use disorder in reward circuitry? Genes Brain Behav 20, e12689 (2021). https://doi.org:10.1111/gbb.12689

43 Palmer, R. H. C. et al. Integration of evidence across human and model organism studies: A meeting report. Genes Brain Behav, e12738 (2021). https://doi.org:10.1111/gbb.12738

44 Binh Tran, T. D. et al. Microbial glutamate metabolism predicts intravenous cocaine self-administration in diversity outbred mice. Neuropharmacology 226, 109409 (2023). https://doi.org:10.1016/j.neuropharm.2022.109409

45 Chesler, E. J. et al. Diversity Outbred Mice at 21: Maintaining Allelic Variation in the Face of Selection. G3 (Bethesda) 6, 3893–3902 (2016). https://doi.org:10.1534/g3.116.035527

46 Sathyanesan, M. et al. A molecular characterization of the choroid plexus and stress-induced gene regulation. Transl Psychiatry 2, e139 (2012). https://doi.org:10.1038/tp.2012.64

47 Morgan, A. P. et al. The Mouse Universal Genotyping Array: From Substrains to Subspecies. G3 (Bethesda) 6, 263–279 (2015). https://doi.org:10.1534/g3.115.022087

48 Broman, K. W., Gatti, D. M., Svenson, K. L., Sen, S. & Churchill, G. A. Cleaning Genotype Data from Diversity Outbred Mice. G3 (Bethesda) 9, 1571–1579 (2019). https://doi.org:10.1534/g3.119.400165

49 Broman, K. W. et al. R/qtl2: Software for Mapping Quantitative Trait Loci with High-Dimensional Data and Multiparent Populations. Genetics 211, 495–502 (2019). https://doi.org:10.1534/genetics.118.301595

50 Gatti, D. M. et al. Quantitative trait locus mapping methods for diversity outbred mice. G3 (Bethesda) 4, 1623–1633 (2014). https://doi.org:10.1534/g3.114.013748

51 Churchill, G. A. Data Structures in QTL Viewer https://github.com/churchill-lab/qtl-viewer/blob/master/docs/QTLViewerDataStructures.md. (2017).

52 Vincent, M. et al. QTLViewer: an interactive webtool for genetic analysis in the Collaborative Cross and Diversity Outbred mouse populations. G3 (Bethesda) 12 (2022). https://doi.org:10.1093/g3journal/jkac146

53 Langfelder, P. & Horvath, S. WGCNA: an R package for weighted correlation network analysis. BMC Bioinformatics 9, 559 (2008). https://doi.org:10.1186/1471-2105-9-559

54 Chesler, E. J. & Langston, M. A. in Proceedings of the 2005 joint annual satellite conference on Systems biology and regulatory genomics 150–165 (Springer-Verlag, San Diego, CA, USA, 2005).

55 Baker, E., Bubier, J. A., Reynolds, T., Langston, M. A. & Chesler, E. J. GeneWeaver: data driven alignment of cross-species genomics in biology and disease. Nucleic Acids Res 44, D555–559 (2016). https://doi.org:10.1093/nar/gkv1329

56 Skelly, D. A., Raghupathy, N., Robledo, R. F., Graber, J. H. & Chesler, E. J. Reference Trait Analysis Reveals Correlations Between Gene Expression and Quantitative Traits in Disjoint Samples. Genetics 212, 919–929 (2019). https://doi.org:10.1534/genetics.118.301865

